# Miltefosine attenuates inflammation, reduces atherosclerosis, and alters gut microbiota in hyperlipidemic mice

**DOI:** 10.1101/2022.08.22.504848

**Authors:** C. Alicia Traughber, Amanda J Iacano, Mariam R Khan, Kalash Neupane, Emmanuel Opoku, Tina Nunn, Naseer Sangwan, Stanley L Hazen, Jonathan D Smith, Kailash Gulshan

**Affiliations:** Center for Gene Regulation in Health and Disease, Cleveland State University, Cleveland, OH 44115, USA; Department of Biology, Geology, and Environmental Sciences, Cleveland State University, Cleveland, OH 44115, USA; Department of Cardiovascular and Metabolic Sciences, Lerner Research Institute, Cleveland Clinic, Cleveland, OH 44195, USA; Center for Microbiome and Human Health, Cleveland Clinic, Cleveland OH 44195, USA

**Keywords:** Atherosclerosis, Cholesterol, Miltefosine, NLRP3 inflammasome, IL-1β, Gut microbiota

## Abstract

Excess cholesterol induces foam cell formation, NLRP3 inflammasome activation, and IL-1β release in atherosclerotic plaques. We have shown previously that Miltefosine increased cholesterol release and dampened NLRP3 inflammasome assembly in macrophages. Here, we show that Miltefosine reduced LPS-induced choline uptake by macrophages and attenuated NLRP3 inflammasome assembly in mice. Miltefosine-fed mice showed reduced plasma IL-1β in a polymicrobial cecal slurry injection model of systemic inflammation. Miltefosine-fed mice showed increased reverse cholesterol transport from macrophages to plasma, liver, and feces. Hyperlipidemic apoE^−/−^ mice fed with Miltefosine showed significantly reduced weight gain and markedly reduced atherosclerotic lesions vs. control mice. 16S rDNA sequencing and analysis showed alterations in the gut microbiota profile of Miltefosine-fed hyperlipidemic apoE^−/−^ vs. control mice, with the most notable changes in *Romboutsia* and *Bacteroidetes species*. Taken together, these data indicate that Miltefosine causes pleiotropic effects on lipid metabolism, inflammasome activity, atherosclerosis, and the gut microbiota.

## Introduction

*Leishmania* parasite preferentially infects phagocytic cells in human hosts, such as macrophages and dendritic cells^1,2^. Macrophages induce NLRP3 inflammasome assembly during *Leishmania* infection in both mice and humans, leading to the processing and release of mature interleukin-1 beta (IL-1β) at the infection site ^3–6^. IL-1β promotes disease progression, as the mice lacking NLRP3, ASC, or caspase 1 showed defective IL-1β production at the infection site and were resistant to cutaneous *Leishmania* infection^4^. In addition, the IL-1β levels in patients with cutaneous leishmaniasis positively correlate with areas of necrosis^7^. Miltefosine is an FDA-approved drug to treat visceral and cutaneous leishmaniasis^8^. The mechanism of action of Miltefosine is not fully clear, but it can freely integrate into the cell membrane and redistribute in the ER, golgi, and mitochondria^8^. Studies from our lab and others have shown that Miltefosine causes cholesterol release from cells^9,10^ and modulates inflammatory responses in a variety of immune cells including macrophages, mast cells, and eosinophils^10–12^. Accumulation and oxidation of excess cholesterol in the arterial intima is the major cause of coronary artery disease (CAD). Oxidized low-density lipoprotein (LDL)-cholesterol (LDL-C) in the artery wall promotes the recruitment of monocytes, which transform into arterial wall macrophages and uptake LDL-C to form lipid-laden foam cells ^13–16^. The oxidized LDL acts as potent activators of the toll-like receptor (TLR) pathway and cholesterol crystals in plaques promote the assembly of NLRP3 inflammasome ^14,17^. The inflammasome-mediated processing of IL-1β may lead to beneficial antimicrobial activity but can also result in amplification of an inflammatory cascade, worsening the pathogenesis of various chronic inflammatory diseases such as metabolic syndrome and atherosclerosis^18–20^. Inflammasome-mediated activation of caspase 1 and caspase 11 also results in cleavage of Gasdermin D (GsdmD), a downstream effector of inflammasome activity required for efficient release of IL-1β from cells^21–26^. Studies from our lab and others have shown the involvement of GsdmD pathway in promoting atherosclerosis in mice and humans^27–31^. In addition to chronic inflammation caused by a disruption in sterol homeostasis, metabolites of the dietary phosphatidylcholine (PC), such as choline and trimethylamine N-oxide (TMAO), can also promote chronic inflammatory pathologies such as atherosclerosis and cardiovascular disease (CVD) in a gut-microbiota dependent manner^32–34^.

In contrast to activated atherogenic pathways, the atheroprotective pathways such as autophagy and reverse cholesterol transport (RCT) are known to become dysfunctional with aging and in advanced atherosclerosis^17,35,36^. We have shown before that Miltefosine promoted cholesterol efflux from macrophages, induced autophagy/mitophagy, and blunted NLRP3 inflammasome assembly. Here, we used a wild-type (WT) and a hyperlipidemic mouse model of atherosclerosis to test the effects of Miltefosine on inflammasome activity, reverse cholesterol transport, atherosclerosis, and gut microbiota.

## Results

### Miltefosine dampened inflammasome activity in human macrophages

The THP-1 monocytes stably expressing ASC-GFP construct were differentiated into macrophages by PMA treatment. The THP-1 macrophages were pretreated with ± 7.5 μM Miltefosine for 16h, followed by priming with LPS for 4h. NLRP3 inflammasome assembly was induced by incubation with 5μM Nigericin for 1h. The THP-1 macrophages treated with LPS and Nigericin formed ASC specks in ~35% of cells, while only 17% of cells pretreated with Miltefosine formed ASC specks (n~ 500 cells for each condition) with p<0.0001 by two-tailed Fisher’s exact test (***Fig.1A, 1B)***. The release of IL-1β from THP-1 macrophages pretreated with Miltefosine was also significantly lower with ~40% reduction vs. control cells (***Fig. 1C***). To ensure that Miltefosine did not affect the priming of macrophages, western blot analysis of pro-IL-1β was performed. Incubation with LPS induced levels of pro-IL-1β equally in both control and Miltefosine-treated cells (***Fig. 1D***). These data indicate that Miltefosine blocks NLRP3 assembly without affecting the LPS mediated priming of macrophages.

**Figure 1:**
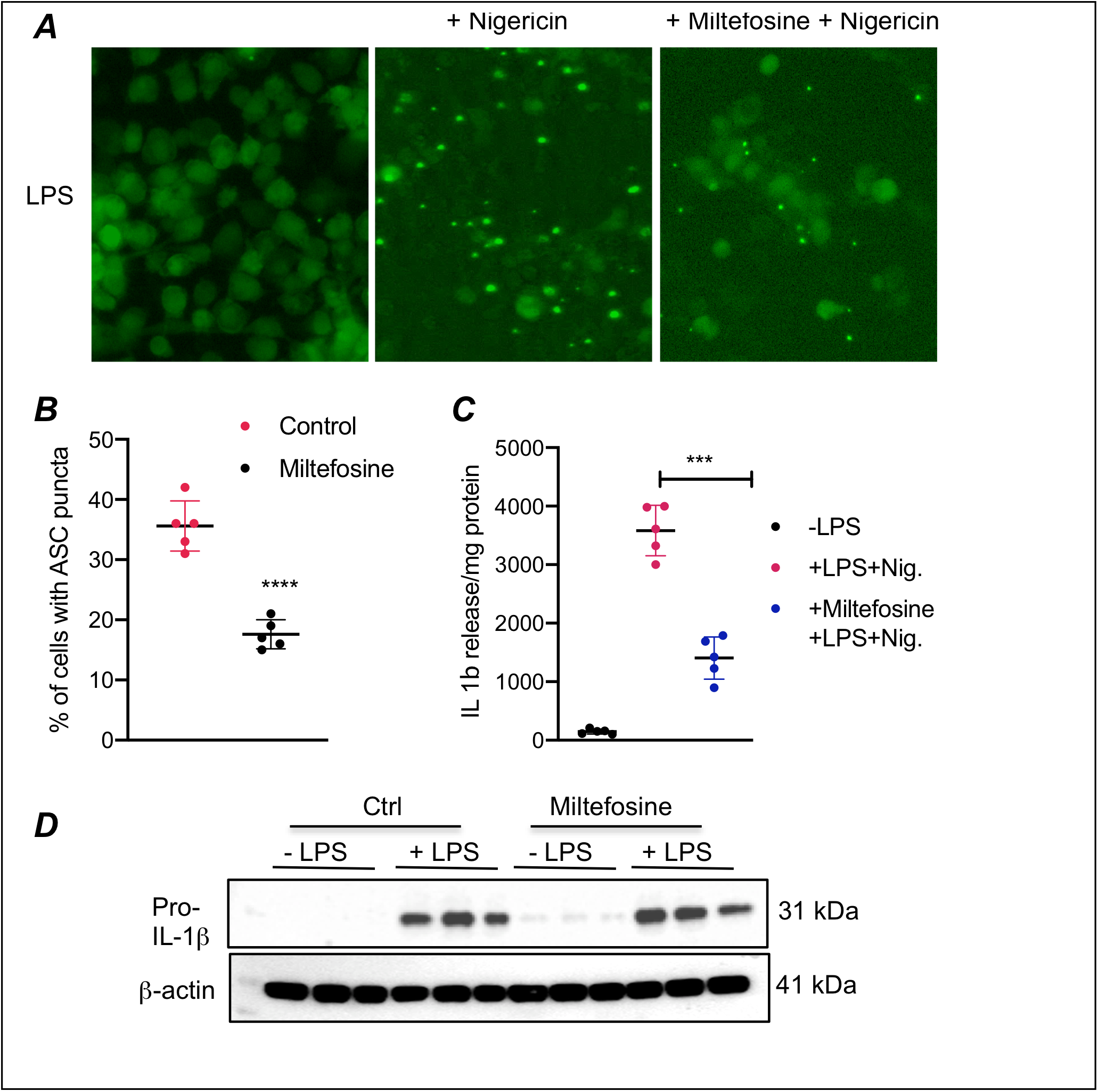
Miltefosine dampened inflammasome assembly in THP-1 macrophages. ***A)*** THP-1 macrophages stably expressing GFP tagged ASC construct were pretreated with ± 7.5 μM Miltefosine, primed by incubation with 1μg/ml LPS and treated with ± 5 mM Nigericin for 1h to induce NLRP3 inflammasome assembly. ASC puncta formation (a marker for NLRP3 inflammasome assembly) was visualized using fluorescent microscopy. ***B)*** The number of ASC puncta in control and Miltefosine treated cells were plotted using separate areas from five wells (with ~ 100 DAPI+ cells) counted (mean ± SD ****, p<0.0001 by two-tailed Fisher exact test using the number of puncta positive and negative cells). ***C)*** ELISA showing the levels of IL-1β in media from THP-1 macrophages treated with ± 7.5 μM Miltefosine ± 1μg/ml LPS and ± 5 mM Nigericin (N=5, mean ± SD ***, p<0.001 by ANOVA, each sample compared with other via using Tukey’s multiple comparisons test. ***D)*** Western blot analysis of LPS induced expression of pro IL-1β in control and Miltefosine treated cells, β-actin was used as loading control.

### Miltefosine blunts endotoxin-induced choline uptake

Previous studies have shown that exposure to LPS induces choline uptake by macrophages to activate the NLRP3 inflammasome while blocking choline uptake prevents NLRP3 inflammasome assembly^37–39^. We tested if one of the mechanisms of Miltefosine anti-inflammasome activity is *via* reduced choline uptake. Bone marrow-derived macrophages (BMDMs) from WT-C57BL6J mice were pretreated with ± 7.5 μM Miltefosine for 16h. The control and Miltefosine treated BMDMs were primed with 1 μg/ml LPS for 4h, followed by incubation with radiolabeled choline at final concentration of 2.5 μCi/ml for 0, 20, or 40 minutes. The extracellular media containing radiolabeled choline was removed after the indicated time and the intracellular radioactivity was determined by liquid scintillation counting. As shown in ***Fig. 2***, there was a significant (~ 35%) reduction in choline uptake in Miltefosine treated vs. control cells at 20 minutes. At 40 minutes, Miltefosine treated cells showed a significant (~ 40 %) reduction in choline uptake vs. control cells. These data indicate that Miltefosine may be blunting inflammasome assembly *via* dampening LPS-induced choline uptake by macrophages.

**Figure 2.**
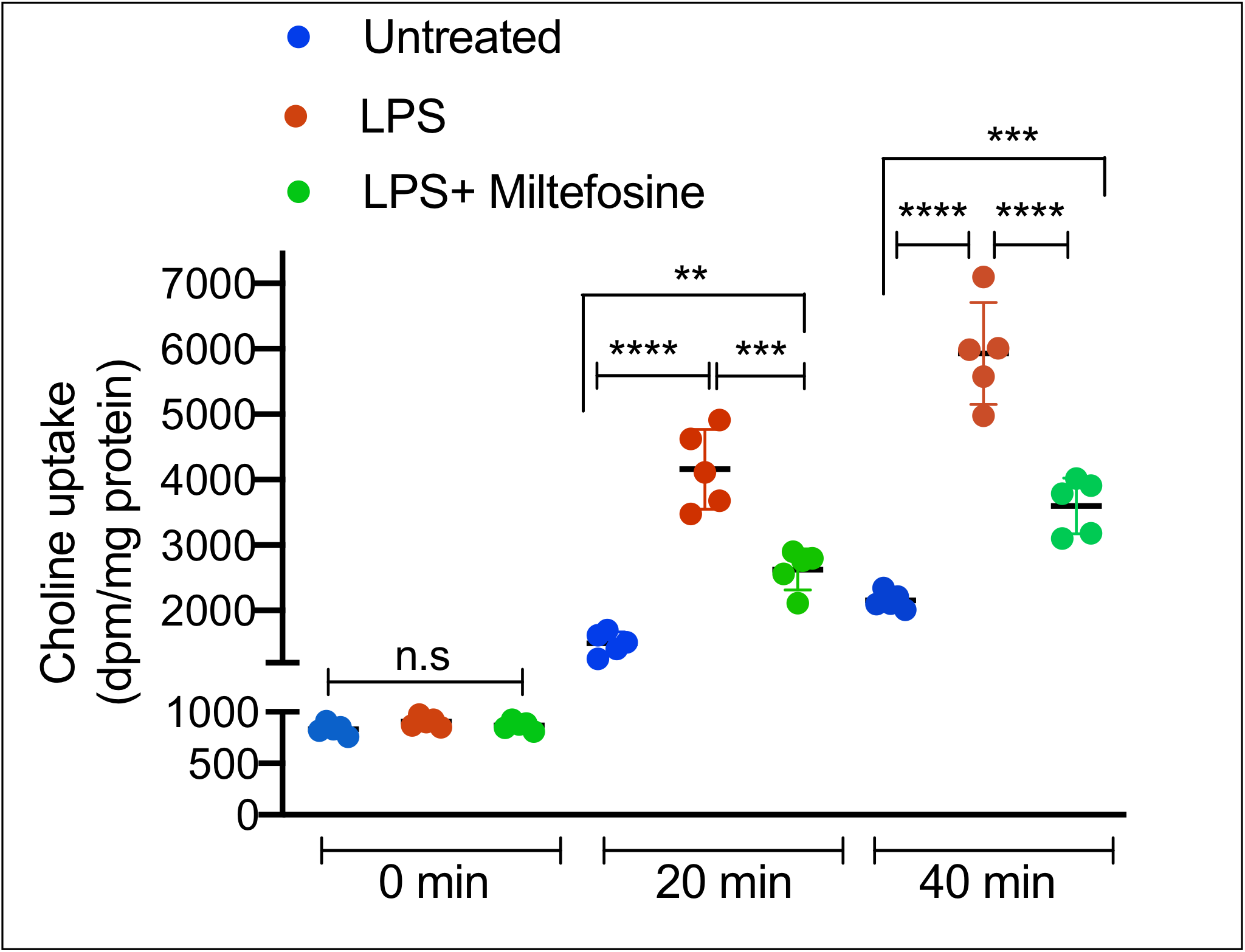
Miltefosine dampened LPS-induced choline uptake. Mouse BMDMs were plated in 6-well plate and treated with ± 7.5 μM Miltefosine for 16h. The cells were primed by incubation with ± 1 mg/ml LPS for 4h, followed by incubation with tritium labeled choline at final concentration of 2.5 μCi/ml ^3^H-choline. The radioactivity uptake assay was performed at 37°C and radioactive dpm counts were determined by using scintillation counter. Each sample in a group was compared with others by ANOVA using Tukey’s multiple comparisons test (N=5, mean ± SD for all groups; for 0 min group, n.s (non significant), for 20 min group, ****, p<0.0001 for untreated vs. LPS treated, ***, p = 0.0002 for LPS vs. LPS + Miltefosine, and **, p = 0.0024 for untreated vs. LPS + Miltefosine, for 40 min group, ****, p<0.0001 for untreated vs. LPS treated, ****, p<0.0001 for LPS vs. LPS + Miltefosine, and ***, p = 0.0006 for untreated vs. LPS + Miltefosine).

### Miltefosine attenuates *in vivo* NLRP3 inflammasome activity

To determine the effect of Miltefosine on *in vivo* NLRP3 inflammasome, the WT mice were fed with a chow diet ± 20 mg/kg/day Miltefosine for 3 weeks. The chow-fed and Miltefosine-fed mice were injected i.p. with 5 μg LPS, followed 4h later with i.p. injection of ATP. The mice were sacrificed 30 min later, and the peritoneal cavity lavage was collected and analyzed for IL-1β levels. As shown in ***Fig. 3***, Miltefosine-fed mice showed significantly reduced IL-1β levels in peritoneal lavage fluid.

**Figure. 3.**
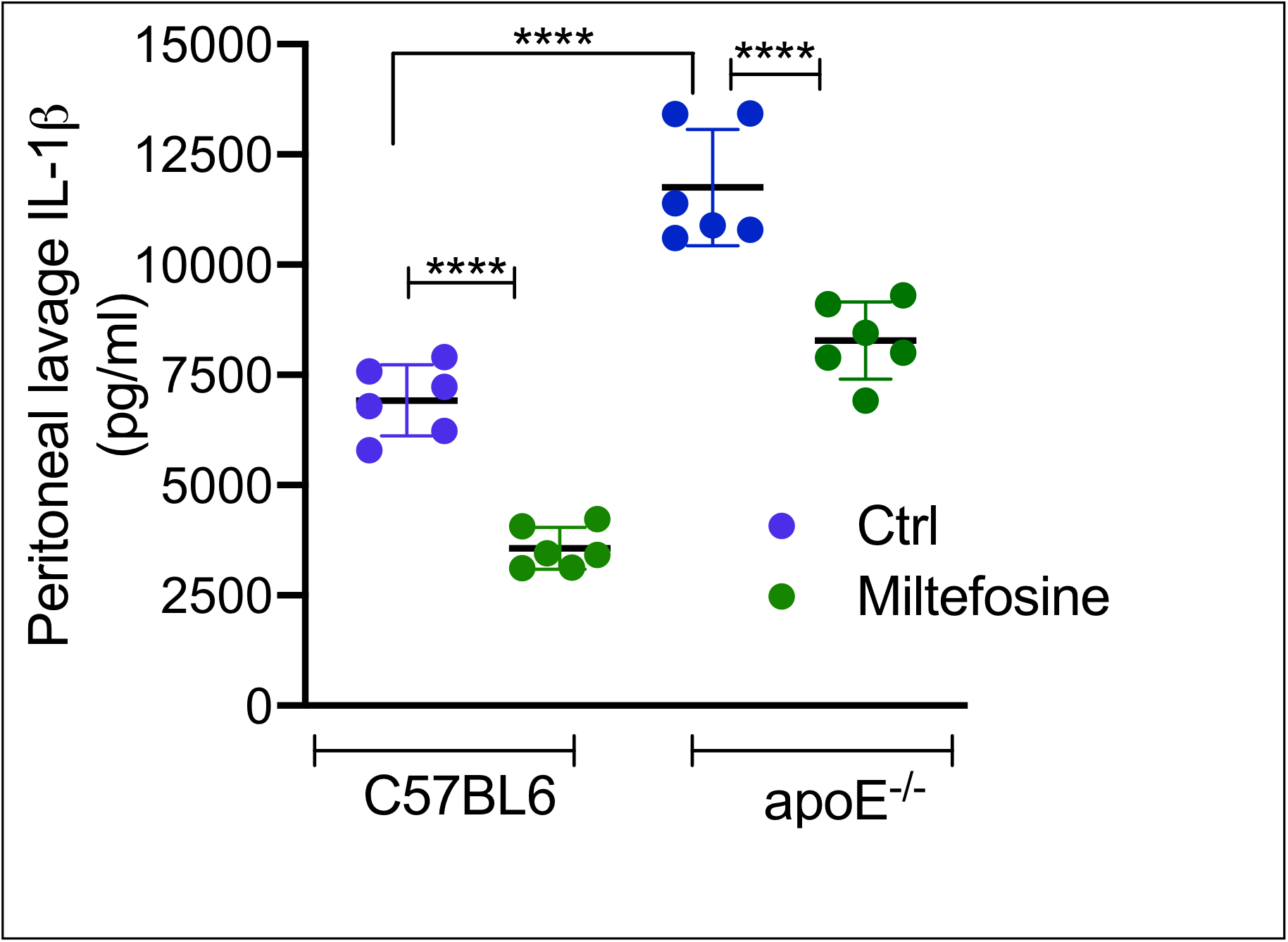
Miltefosine attenuated in vivo NLRP3 inflammasome activity. Age-matched (10-week old) male WTC57BL6J mice fed with chow ± 20 mg/kg Miltefosine for 3 weeks or C57BL6J-apoE^−/−^ knockout mice fed with WTD ± 50 mg/kg Miltefosine for 2 weeks were used. The mice were primed for inflammasome assembly by an I.P. injection of LPS (5mg/mouse). After 4h of LPS injection, the NLRP3 inflammasome assembly was induced by I.P. injection of ATP (0.5 ml of 30 mM, pH 7.0). The peritoneal cavity was lavaged with 5 ml sterile PBS, and IL-1β levels in peritoneal lavage were determined by ELISA (N=6, mean ± SD for all groups, **** p <0.0001 for C57BL6J-control vs. C57BL6J-Miltefosine, **** p <0.0001 for C57BL6J-control vs. apoE^−/−^ -control, **** p <0.0001 for apoE^−/−^-control vs. apoE^−/−^ -Miltefosine by two-tailed t-test).

To determine if Miltefosine can also reduce IL-1β levels in hyperlipidemic conditions, the apoE^−/−^ mice fed with a western type diet (WTD) ± 50mg/kg/day Miltefosine for 3 weeks were injected with LPS +ATP for inflammasome induction. As shown in ***Fig. 3***, the WTD-fed apoE^−/−^ mice showed robust IL-1β levels in peritoneal lavage, while Miltefosine-fed mice showed ~ 65% reduction. Control mice received either saline or LPS + saline injections and showed negligible levels of IL-1β in peritoneal lavage (***Fig. S1***). These data indicated that Miltefosine dampened *in vivo* NLRP3 inflammasome activity in mice.

### Miltefosine reduced inflammation-induced cytokine release in hyperlipidemic mice

Hyperlipidemia and sepsis-induced inflammation promote atherosclerosis and CVD in humans^40,41,42^. We used WT-C57BL6J and and C57BL6J-apoE^−/−^ mice for the studies. The tested different Miltefosine doses in a 6 week pilot studies with WT mice and 20mg/kg/day for chow diet and 50mg/kg/day for western type diet were chosen based on no weight loss or absence of any other visible adverse effect. To determine the effect of Miltefosine treatment on inflammatory responses in hyperlipidemic animals, a polymicrobial cecal slurry injection model of peritonitis was used^43^. The cecal slurry injection dose of 4μl/g body weight was selected as this dose did not induce mortality but still led to a transient decrease in body temperature to 35°C. The apoE^−/−^ mice fed with a WTD diet ± 50 mg/kg/day Miltefosine for 3 weeks were i.p. injected with cecal slurry, and the plasma was collected at 2, 4, and 6h post-injection. As shown in ***Fig. 4***, the Miltefosine-fed mice group had significantly reduced IL-1β release in plasma vs. control mice. These data indicated that Miltefosine reduce inflammatory responses in hyperlipidemic mice.

**Figure. 4.**
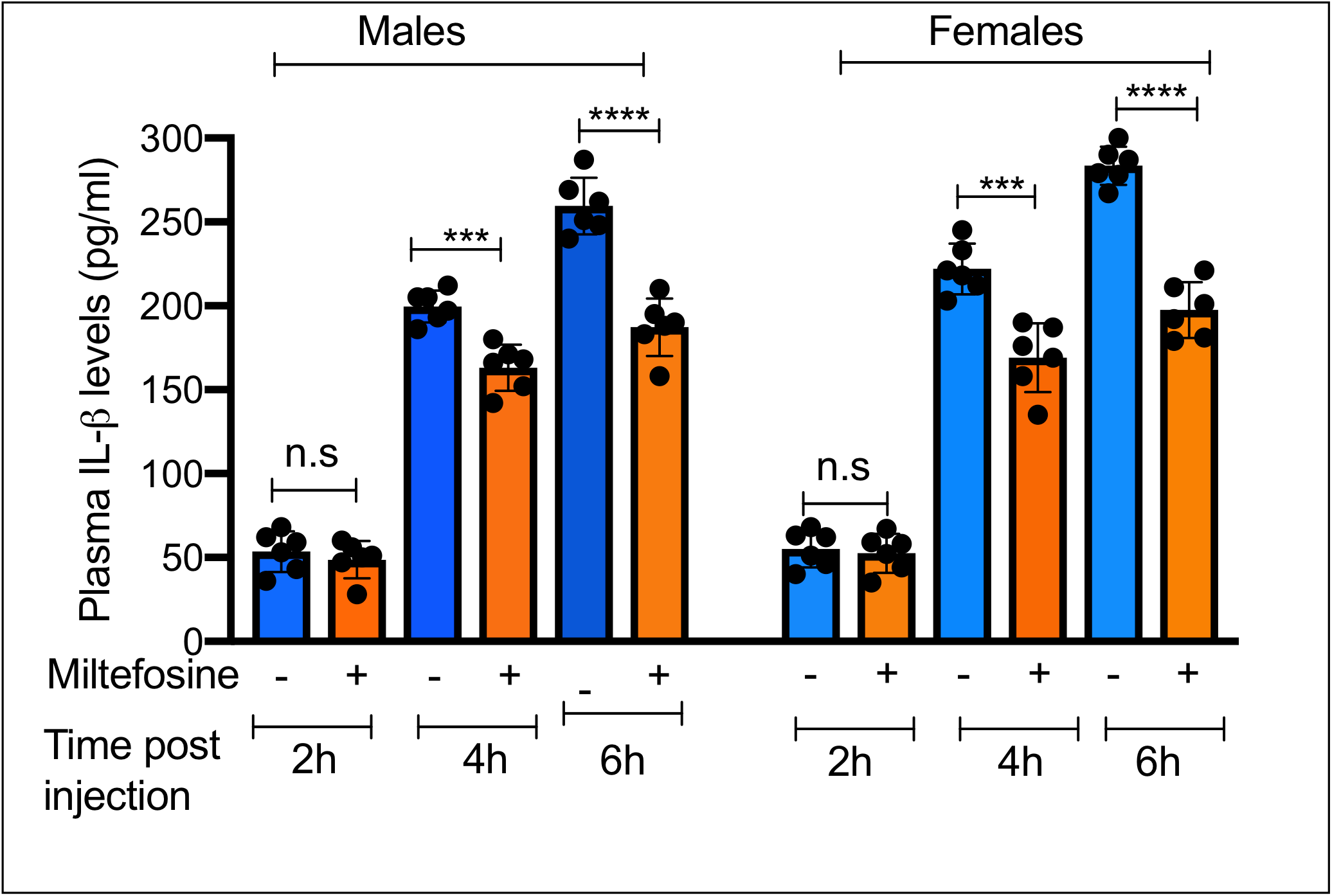
Miltefosine dampened IL-1β release in polymicrobial cecal slurry mouse model of systemic inflammation. The age and sex-matched apoE^−/−^ mice were fed with WTD or WTD + 50 mg/kg Miltefosine for 3 weeks. The mice were injected i.p with cecal slurry (from WT C57BL6J mice) with the dose of 4 μl/gram of body weight. The blood was collected by tail bleed at indicated times after injection and plasma IL-1β levels were determined by ELISA (N=6, mean ± SD for all groups; for 2 h males and females group, n.s (non significant), for 4h male group ***, p=0.0003, for 4 h female group ***, p=0.0005, for 6h male group ****, p<0.0001, for 6 h female group, ****, p<0.0001 with two-tailed t-test.

### Miltefosine increases reverse cholesterol transport in mice

To determine if Miltefosine treatment in mice leads to increased reverse cholesterol transport (RCT), the RCT assay^44^ was performed in WT mice, as apoE^−/−^ mice are not suitable for these studies due to inherent defects in lipoprotein packaging and excessive plasma cholesterol pool even with chow-diet feeding. Donor BMDMs from WT mice were loaded with 100 μg/ml acetylated-low-density lipoprotein (Ac-LDL) and radiolabeled ^3^H-cholesterol for 48 h to generate foam cells, which were implanted s.c into the back flanks of he 9-week old WT recipient mice, who were then fed chow diet or chow diet containing 20 mg/kg/day Miltefosine. The radioactive cholesterol mobilized to plasma was determined by collection of blood samples at 24 h, 48 h, and 72 h. The liver and feces samples were collected post euthanasia and processed to determine the percent of RCT to these pools. As shown in ***Fig. 5A***, Miltefosine increased RCT to plasma in mice by ~30% at 24 h, by 26% at 48 h, and by ~20% at 72 h, RCT to feces showed ~ 31% increase at 24 h, 26% increase at 48 h, and ~51% increase at 72 h in Miltefosine-fed vs. chow-fed mice (***Fig. 5B***). RCT to the liver was also increased by ~21% in Miltefosine-fed mice (***Fig. 5C***). These data indicated that Miltefosine increased RCT *in vivo*.

**Figure. 5.**
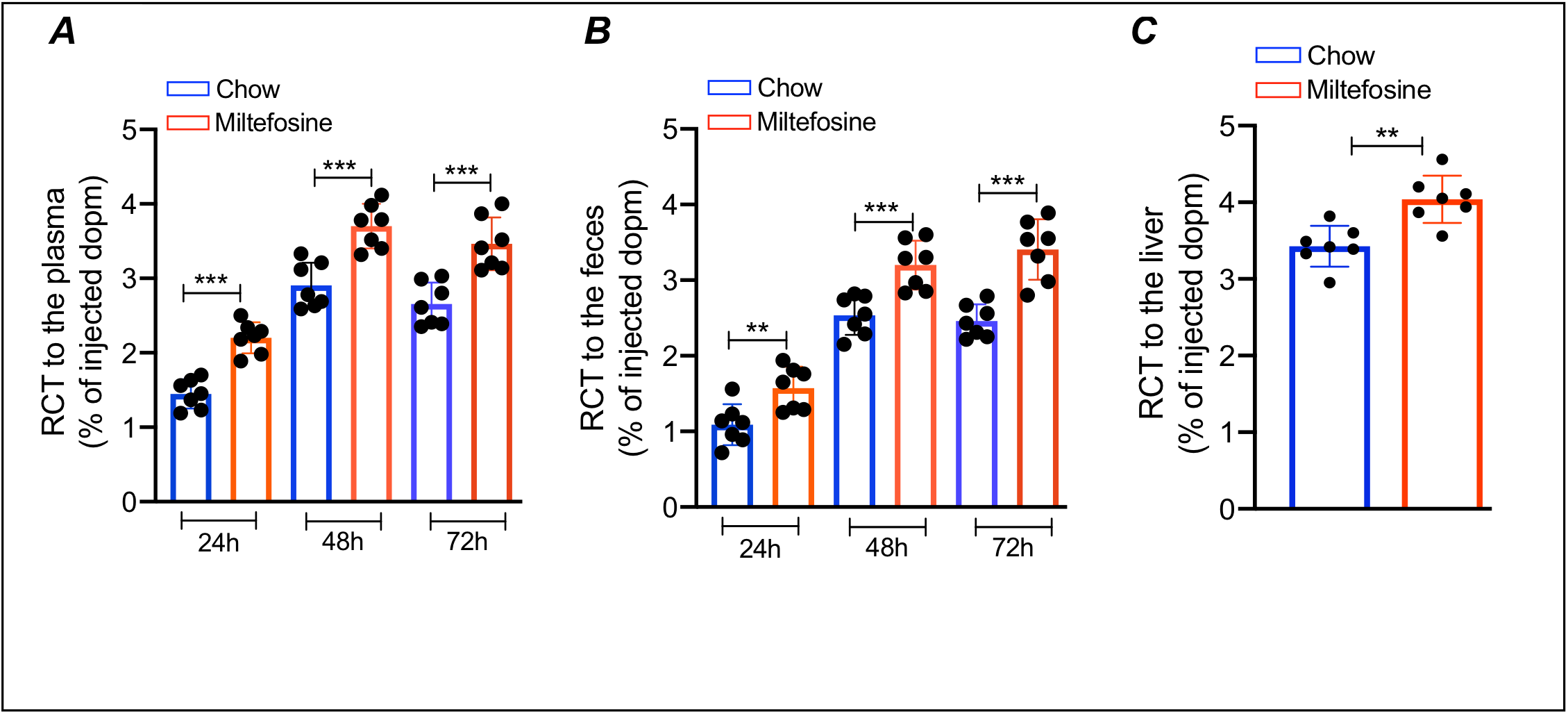
Miltefosine increased RCT in mice. The foam cells prepared by loading of BMDMs isolated from WT C57BL6J mice with ^3^H-labeled cholesterol, were transplanted into age-matched WT male recipient mice fed with either chow or chow+ 20 mg/kg Miltefosine for 3 weeks. ***A)*** RCT to plasma determined at 24, 48, and 72 h (N=7, mean ± SD for all groups, ***, p = 0.004 to 0.005 by two-tailed t-test). ***B)*** RCT to feces determined at 24, 48, and 72 h (N=7, mean ± SD for all groups, for 24 h group **, p = 0.0067 for chow vs. Miltefosine by two-tailed t-test, for 48h group, **, p = 0.0012, and for 72 h group, ***, p=0.0001 for chow vs. Miltefosine by two-tailed post t-test. ***C)*** RCT to liver determined at 72 h (N=7, mean ± SD, **, p = 0.0018 for chow vs. Miltefosine by two-tailed t-test.

### Miltefosine reduced atherosclerosis in hyperlipidemic mice

To determine if Miltefosine can reduce atherosclerosis, 6-week old apoE^−/−^ mice were weaned onto the atherogenic western type diet (WTD) or WTD containing 50mg/kg/day Miltefosine for 9 weeks. The Miltefosine-fed mice gained significantly less body weight vs. the WTD-fed group (***Fig. 6A***). No significant differences were found in food intake in WTD vs. WTD + Miltefosine group (***Fig. 6B)***. The oil red O positive aortic root atherosclerotic lesion areas were significantly reduced in Miltefosine-fed group with ~ 50% reduction in atherosclerotic lesions in both males and females (***Fig. 6C, 6D***). There was a trend toward lowered total cholesterol and higher HDL-cholesterol in the plasma of the Miltefosine-fed group, but the changes did not achieve levels of significance (***Fig. 6E)***. These data indicated that Miltefosine reduces atherosclerosis and this effect may not be completely dependent on the reduction in cholesterol load.

**Figure. 6.**
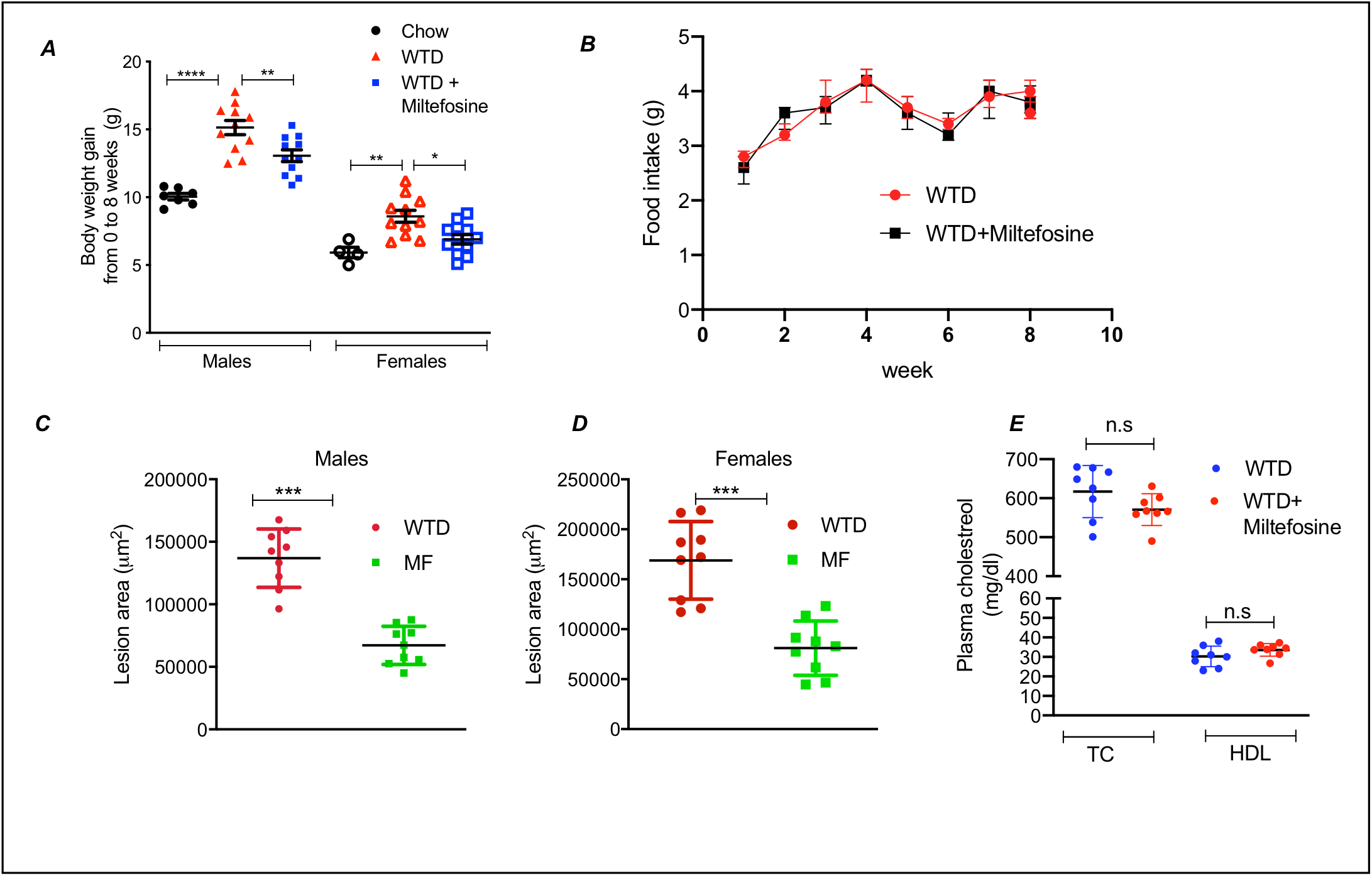
Miltefosine reduced atherosclerosis in hyperlipidemic mice. The 5-week old apoE^−/−^ mice from both sexes were treated with either WTD or WTD + 50 mg/kg Miltefosine diet for 9 weeks. The control apoE^−/−^ mice were fed chow diet throughout the course of the study. ***A)*** The body weight (BW) gain during course of study. The gain in BW was calculated by subtracting BW at end-point (week 14) from BW at beginning of study (week 5) and was plotted, for male mice group, **** indicate p<0.0001 for chow vs. WTD, **** indicate p<0.0001 for chow vs. WTD + Miltefosine, and *** indicate p = 0.0003 for WTD vs. WTD + Miltefosine with ANOVA. ***B***) Average food intake in mice fed with WTD or WTD + 50 mg/kg Miltefosine. The measurements were taken weekly till the end of the study, p= n.s (non-significant). ***C***) Quantification of aortic root lesions in male mice fed with either WTD or WTD + 50mg/kg Miltefosine. N=12, *** indicate p <0.0005, by two-tailed t-test). ***D***) Quantification of aortic root lesions in female mice fed with either WTD or WTD + 50mg/kg Miltefosine, N=12, *** indicate p <0.0005, by two-tailed t-test). ***E***) Plasma cholesterol (total and HDL) in apoE^−/−^ mice fed with WTD or WTD + Miltefosine diet was determined by Stan-bio kit following manufacturer’s instruction (N=8 for males and females, n.s= non-significant by two-tailed t-test).

### Altered gut microbiota in Miltefosine treated mice

Gut microbiota play a major role in the progression of atherosclerosis and CVD^32,45,46^. Miltefosine was originally identified as an anti-cancer compound but was later shown to be effective against a variety of microbes including bacteria^8^. Thus, we determined if Miltefosine modulates the gut microbiota composition by performing a 16s ribosomal DNA (16S rDNA)-based qPCR sequencing. Fresh fecal samples were collected from apoE^−/−^ mice fed for 9 weeks with either chow diet, WTD, or WTD + Miltefosine. The DNA isolated from these samples was analyzed for the microbial species profile and alpha and beta diversity across different groups. Alpha diversity is a measure of microbiome diversity/complexity in each sample, while the beta diversity is a measure of similarity or dissimilarity between groups. The gut microbiota profile was significantly different in three groups with alterations in alpha and beta diversity in chow vs. other groups and WTD-fed vs. WTD + Miltefosine-fed group (***Fig. 7A, 7B***). The WTD-fed mice showed a marked increase in *Romboutsia* species vs. chow-fed mice (***Fig. 7C)*** and this effect was blunted in Miltefosine-treated mice. The Miltefosine-treated group showed increased levels of *Bacteroides* species vs. the WTD-fed group (***Fig. 7C***). These data indicate that Miltefosine alters the gut microbiota profile and this alteration could serve as one of the mechanisms for Miltefosine’s anti-atherosclerotic property.

**Figure. 7.**
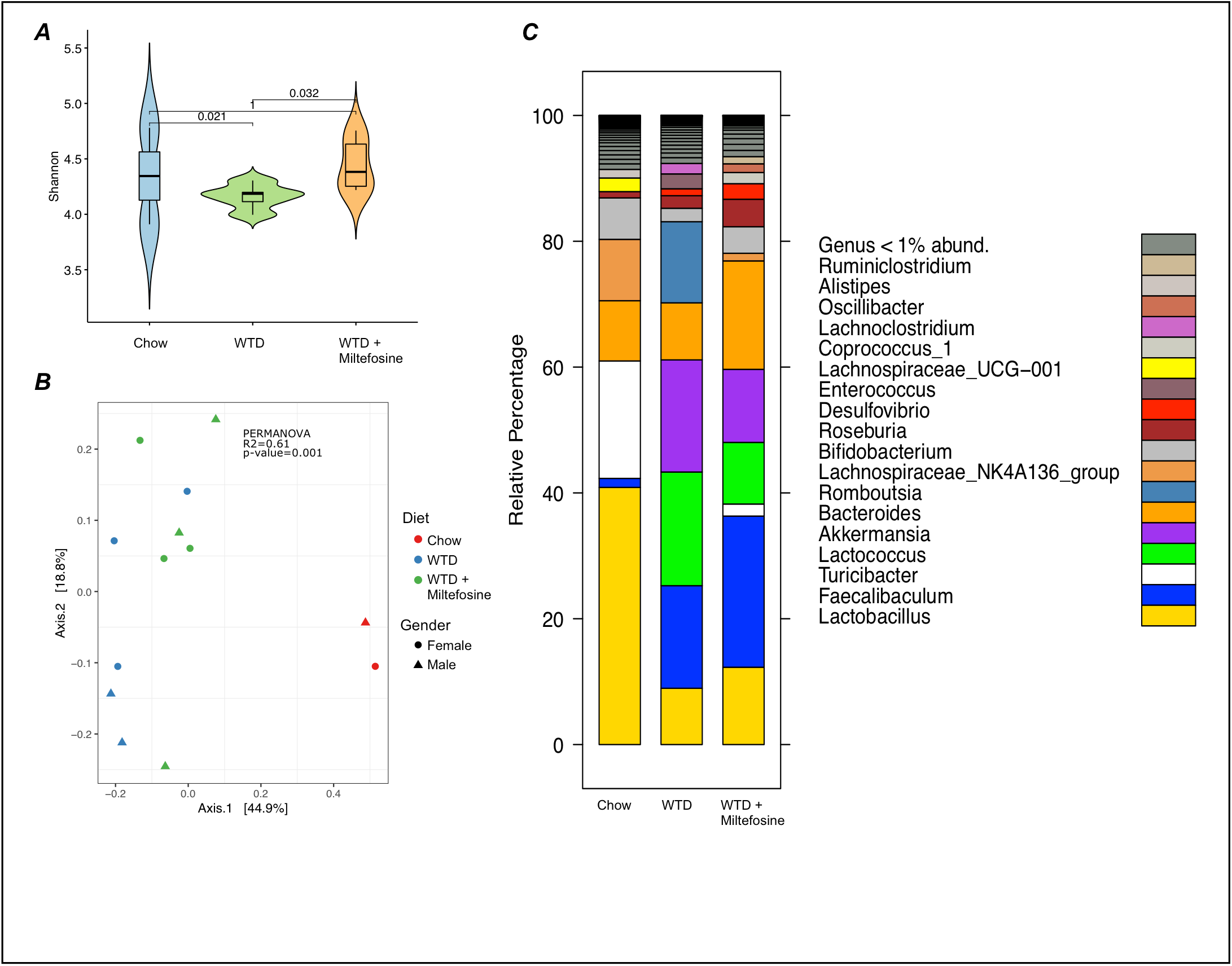
Miltefosine alters gut microbiota profile in hyperlipidemic mice. The 5-week old apoE^−/−^ mice were fed with either WTD or WTD + 50 mg/kg Miltefosine diet for 9 weeks. The control apoE^−/−^ mice were fed chow diet throughout. Fresh feces were collected and 16s rDNA sequencing was performed. ***A)*** Alpha diversity in gut microbial profile of mice fed with different diets. ***B)*** Beta diversity of microbiota in gut microbial profile of mice fed with different diets. ***C)*** Total diversity in gut microbial profile of mice fed with different diets.

## Discussion

Miltefosine is an alkyl-lysophospholipid analog with *in vitro* activity against various *Leishmania* species. The activity of NLRP3 inflammasome and release of IL-1β during leishmaniasi*s* had been reported in mouse models as well as in human patients^3–6^. The activation of the NLRP3 inflammasome, instead of serving as a tool to clear the parasitic infection, was found to be detrimental during leishmaniasis and the mice lacking NLRP3, ASC, or caspase 1 were shown to be resistant to cutaneous infection^4^. Furthermore, the amount of IL-1β positively correlates with areas of necrosis in cutaneous leishmaniasis patients^7^. We found that Miltefosine reduced *in vivo* NLRP3 inflammasome assembly and IL-1β release in human macrophages (***Fig. 1***). Thus, in addition to killing the pathogen, Miltefosine may also be modulating host immune responses to *Leishmania* infection by regulating inflammasome activity.

Miltefosine displays a range of activities such as anticancer, antimicrobial, effects on lipid metabolism homeostasis, and immune cell function ^8,12,47,48^, but the exact mechanism of action is not fully understood and its activities seem to be cell-type dependent. Miltefosine altered phosphatidylserine (PS) and phosphatidylinositol 4,5-bisphosphate (PIP2) localization in macrophages^10^, and negatively affect phosphatidylcholine (PC) and sphingolipid biosynthesis^48,49^. Previous studies have shown that the uptake of PC biosynthetic precursors, such as choline, precedes NLRP3 inflammasome assembly and IL-1β release^37,38^. The cells exposed to LPS stimuli up-regulate the expression of choline transporter to import more choline for generating PC. We found that Miltefosine reduced choline uptake in LPS-induced macrophages (***Fig. 2***). Thus, Miltefosine may be dampening inflammasome assembly due to combinatorial inhibition of PC biosynthesis and choline uptake.

We found that Miltefosine reduced LPS-induced *in vivo* NLRP3 inflammasome activity and IL-1β release (***Fig. 3, 4***). Priming of the inflammasome pathway by LPS is dependent on TLR receptors, which are enriched in membrane lipid rafts. Miltefosine-mediated disruption of lipid rafts can attenuate TLR signaling pathway^10,50^, thus we tested the effects of Miltefosine on direct NLRP3 inflammasome activity in live animals. We chose the peritoneal cavity as the site for NLRP3 inflammasome activity as it contains the liver, spleen, GI tract, and a variety of immune cells. The high percentage of naïve tissue-resident macrophages in the peritoneal cavity also makes it a suitable site for testing *in vivo* NLRP3 inflammasome activity. We found that mice fed with Miltefosine had lower IL-1β levels in peritoneal lavage as well as in plasma. Miltefosine, thus may not only be involved in clearing leishmanial infection in human hosts by directly killing the pathogen but also by dampening the detrimental NLRP3-IL-1β pathway to prevent tissue injury.

The limitation of the *in vivo* inflammasome study is that the individual contribution of B cells, T cells, or macrophages in Miltefosine-mediated blockage of NLRP3 activity is not dissected. Previous studies showed that Miltefosine promoted cholesterol removal from the cells^9,10^, but the effects of Miltefosine on cholesterol efflux in live animals are not clear. We found that Miltefosine-fed mice showed increased removal of cholesterol from the transplanted foam cells (***Fig 5***). We speculated that the *in vivo* cholesterol-removing activity of Miltefosine may be used to treat metabolic diseases caused by hyperlipidemia. We tested if Miltefosine can reduce atherosclerosis, a disease promoted by hyperlipidemia and inflammation. The apoE^−/−^ mice were used, as these mice are prone to severe atherosclerosis upon feeding with a cholesterol-rich WTD diet. The apoE^−/−^ mice fed with WTD +Miltefosine showed a significant reduction in atherosclerotic plaque formation vs. mice fed with WTD alone (***Fig 6***).

The chemical structure of Miltefosine is similar to lyso-PC^8^ and Miltefosine negatively affects PC biosynthesis^8^. Dietary PC is metabolized by gut microbiota and converted to choline and atherogenic metabolite trimethylamine N-oxide (TMAO)^32,45^. Given the antimicrobial activity of Miltefosine, we tested if Miltefosine can alter the gut microbiota of hyperlipidemic mice. We found that gut microbiota from the mice fed with WTD had lower alpha diversity compared to chow-fed mice, while the mice fed with WTD + Miltefosine showed increased alpha diversity vs. mice fed with WTD. Low alpha diversity of gut microbiota has been observed in several metabolic diseases such as obesity, hyperinsulinemia, and dyslipidemia. The WTD-fed mice showed increased levels of *Enterococcus* vs. chow-fed mice, while Miltefosine feeding reduced levels of *Enterococcus* vs. WTD-fed mice. Miltefosine also increased levels of *Bacteroides* species vs. WTD. We also found a marked alteration in levels of *Romboutsia*, with WTD-fed mice showing an increase vs. chow-fed mice, while mice fed with WTD + Miltefosine showed marked reduction in *Romboutsia* levels vs. WTD-fed mice. Previous studies have shown a differential abundance of *Enterobacteriaceae, Bacteroides, and Romboutsia* in atherosclerosis^51–54^. One of the mechanisms by which Miltefosine can alter gut microbiota is via its anti-bacterial properties^8^. Miltefosine may selectively promote the growth of athero-protective gut microbes while inhibiting the growth of athero-promoting bacterial species.

The limitations of our study are that we did not measure levels of Miltefosine in mouse plasma or determined the tissue-specific distribution or *in vivo* half-life of Miltefosine. There is no straightforward assay to measure Miltefosine in mouse plasma, and the mass-spectrometry methods are not well standardized with reported high variability and only a handful of studies using this method^55^. Studies using radioactive Miltefosine have shown that it has a wide distribution in the body with high levels in the kidney, intestinal mucosa, liver, and spleen with a half-life of > 6 days^8^. We did not provide the exact mechanism through which Miltefosine elicits an anti-atherosclerotic effect or provide direct evidence that Miltefosine effects are mediated via gut microbial processes. These changes may only be associated with the other observed phenotypes of Miltefosine such as reduction in lipids and atherosclerosis. To prove a mechanistic link would require a transplantation of cecal microbes and transmission of Miltefosine-dependent anti-atherosclerotic effects in germ-free (GF) mice. The successful transmission would also only show some of the effects of the Miltefosine, in part, are transmissible with microbial transplantation to GF recipients, but it still would not provide evidence of particular microbial metabolites, such as TMAO, being the sole mediator of Miltefosine activity amongst all the other numerous microbial processes that were transplanted.

Miltefosine is a broad-spectrum antimicrobial agent that was originally developed in the 1980s as an anticancer drug. Given that Miltefosine had been used in humans for decades, it can be potentially used in lower doses as a stand-alone or as an adjuvant to LDL lowering therapies to treat inflammatory diseases such as atherosclerosis. In support of Miltefosine as a potential anti-atherosclerotic molecule, previous studies have shown that alterations in gut microbial species, either via dietary interventions with chemical compounds such as Metformin and resveratrol or via gavage inoculation, can impact the progression of atherosclerosis^51,54,56,57^. Further studies are required to determine the efficacy of Miltefosine for treating inflammatory metabolic diseases in humans.

## Material and Methods

### Cell culture

The THP-1 cells were obtained from ATCC and were cultured in RPMI-1640 medium supplemented with 2-mercaptoethanol to a final concentration of 0.05 mM and fetal bovine serum to a final concentration of 10% + penicillin G sodium (100 U/ml) and streptomycin (100 μg/ml). THP-1 cells were differentiated to macrophages using 100 nM phorbol 12-myristate 13-acetate (PMA, Sigma P8139), the PMA was also included in downstream experimental design. Miltefosine (850337P) was obtained from Avanti Polar. Radioactive ^3^H-cholesterol (NET139001) and ^3^H-choline chloride (NET109001) were obtained from Perkin-Elmer Life Sciences. The antibodies for NLRP3, IL-1β, and β-actin were from Cell Signaling.

### Mice and Diets

All animal experiments were performed in accordance with protocols approved by the Cleveland State University and Cleveland Clinic Institutional Animal Care and Use Committee. The C57BL6J mice were purchased from the Jackson laboratory and the apoE^−/–^-C57BL6J mice were bred in-house. Details of mice maintenance and diets are included in *supplementary material & methods* section.

### Isolation of Bone marrow derived macrophages

The WT C57BL6J mice were maintained on chow diet and sacrificed at 16 weeks of age. Detailed method is included in *supplementary material & methods* section.

### Radioactive choline uptake assay

Choline uptake was determined by measuring intracellular ^3^H-choline chloride (PerkinElmer Life Sciences) in mouse BMDMs over time. Detailed method is included in *supplementary material & methods* section.

### In vivo NLRP3 inflammasome activity

*In vivo* NLRP3 inflammasome assembly was induced by LPS and ATP injections in mice, as described earlier^27^. Detailed method is included in *supplementary material & methods* section.

### Mice RCT assay

RCT assays were performed as described earlier^27^. Detailed method is included in *supplementary material & methods* section.

### Cholesterol measurements

Total cholesterol was measured by using Stan Bio Total cholesterol reagent (#1010-225) and plasma HDL-C was determined using ultracentrifugation and precipitation with HDL precipitation reagent (StanBio # 0599020), following manufacturer’s instructions.

### Atherosclerotic Lesion Quantification

Mice were sacrificed by CO_2_ inhalation and weighed at 21 weeks of age. Whole blood was collected from the retroorbital plexus into a heparinized glass capillary, mixed with EDTA and spun in a microfuge to obtain plasma. The circulatory system was perfused with 10 mL PBS and the heart was excised and preserved in 10% phosphate buffered formalin. Hearts were sectioned using Leica cryostat (CM1860) and sections containing aortic sinus were embedded in OCT medium. Quantitative assessment of atherosclerotic plaque area in the aortic root was performed and lesion areas were quantified as the mean value in multiple sections at 80 μm intervals using Image Pro software (Media Cybernetics).

### 16S rDNA sequencing and analysis

The fresh feces samples were collected from mice using sterile forceps and spatula, and the microbial DNA was extracted using Qiagen DNeasy PowerSoil Pro Kit (47016), following manufacturer’s instructions. The DNA samples were subjected to 16S rRNA gene amplification and sequencing using methods explained earlier^58^. Raw 16S amplicon sequence and metadata, were *demultiplexed using split_libraries_fastq*.py script implemented *in QIIME1.9.1^59^*. Demultiplexed fastq file was split into sample specific fastq files using split_sequence_file_on_sample_ids.py script from Qiime1.9.1^59^. Individual fastq files without non-biological nucleotides were processed using Divisive Amplicon Denoising Algorithm (DADA) pipeline^60^.

### Statistical analyses

Data are shown as mean ± SD. Comparisons of 2 groups were performed by a 2-tailed t test, and comparisons of 3 or more groups were performed by ANOVA with Bonferroni posttest. All statistics were performed using Prism software (GraphPad). For microbiome statistical analysis, the output of the dada2 pipeline (feature table of amplicon sequence variants (an ASV table)) was processed for alpha and beta diversity analysis using *phyloseq^61^*, and microbiomeSeq (http://www.github.com/umerijaz/microbiomeSeq) packages in R. Alpha diversity estimates were measured within group categories using estimate_richness function of the *phyloseq* package^61^. Multidimensional scaling (MDS, also known as principal coordinate analysis; PCoA) was performed using Bray-Curtis dissimilarity matrix^62^ between groups and visualized by using *ggplot2* package^63^. We assessed the statistical significance (*P* < 0.05) throughout and whenever necessary, we adjusted *P*-values for multiple comparisons according to the Benjamini and Hochberg method to control False Discovery Rate^64^ while performing multiple testing on taxa abundance according to sample categories. We performed an analysis of variance (ANOVA) among sample categories while measuring the of α-diversity measures using plot_anova_diversity function in *microbiomeSeq* package (http://www.github.com/umerijaz/microbiomeSeq). Permutational multivariate analysis of variance (PERMANOVA) with 999 permutations was performed on all principal coordinates obtained during PCoA with the *ordination* function of the *microbiomeSeq* package. Linear regression (parametric test), and Wilcoxon (Non-parametric) test were performed on ASVs abundances against meta-data variables levels using their base functions in R^65^.

## Non-standard abbreviations and acronyms

ApoA1: apolipoprotein A1
ApoE: apolipoprotein E
ABCA1: ATP-binding cassette transporter A1
AIM2: absence in melanoma 2
ASC: apoptosis-associated speck-like protein containing a CARD.
BMDMs: bone marrow-derived macrophages
CVD: cardiovascular disease
GsdmD: Gasdermin D
HDL-C: high-density lipoprotein-cholesterol
IL-1β: Interleukin 1-beta
LDL-C: low-density lipoprotein-cholesterol
NLRP3: NOD-like receptor family pyrin domain-containing 3
PC: phosphatidylcholine
PIP2: phosphatidylinositol 4,5-bisphosphate
PS: phosphatidylserine
RCT: reverse cholesterol transport
TLR: toll-like receptor
TMAO: trimethylamine N-oxide
WTD: western type diet

## Acknowledgments

This research was supported by NIH-NHLBI grant R01-158148 and Cleveland State University startup funds to K.G.

