## Supplementary material for "Miltefosine attenuates inflammation, reduces atherosclerosis, and alters gut microbiota in hyperlipidemic mice"

Running title: Miltefosine reduces atherosclerosis in mice.

C. Alicia Traughber[1,2,3,*] , Amanda J Iacano[3,*], Mariam R Khan[1,2], Kalash Neupane[1,2], Emmanuel Opoku[3], Tina Nunn[4], Naseer Sangwan[3,4], Stanley L Hazen[3,4], Jonathan D Smith[3], and Kailash Gulshan*[1,2,3]

### SI Materials and Methods

***Mice maintenance and diets:*** The animals were maintained in a temperature-controlled facility with a 12-h light/dark cycle with free access to food and water. The standard chow diet (SD, 20% kcal protein, 70% kcal carbohydrate and 10% kcal fat, Harlan Teklad) was used for regular maintenance and breeding. For generating hyperlipidemia for atherosclerosis studies, mice were fed an atherogenic Western type diet (WTD) (Envigo, 0.2% cholesterol with 42% adjusted calories from fat, TD.88137). Miltefosine was milled in the standard chow or Western type diet by Envigo.

***Radioactive choline uptake assay***: Choline uptake was determined by measuring intracellular $^3$H-choline chloride (Perkin-Elmer Life Sciences) over time. BMDMs were seeded into 12-well plates and treated with or without 7.5 μM Miltefosine for 16h. Treated and control cells were washed twice with sterile PBS, followed by incubation with Krebs-Ringer-HEPES (KRH) buffer (130mM NaCl,

1.3 mM KCl, 2.2 mM $CaCl_2$, 1.2 mM $MgSO_4$, 1.2 mM $KH_2PO_4$, 10 mM HEPES, pH 7.4 and 10 mM glucose) for 1h. The cells were washed again, followed by the addition of KRH buffer containing 2.5 μCi/ml $^3$H-choline chloride and incubation at 37°C for 20 minutes. Cells were washed twice with ice-cold KRH buffer and were lysed with 0.1M NaOH and the radioactivity was determined by liquid scintillation counting. The uptake was plotted as dpm/mg of protein.

***In vivo NLRP3 inflammasome activity***: Mice were fed with either chow, chow+ Miltefosine, WTD or WTD + Miltefosine for 3 weeks. Mice were I.P. injected with either 5 µg LPS, or sterile PBS. After 4 h of LPS or PBS injection, the NLRP3 inflammasome assembly was induced by I.P. injection of ATP (0.5 ml of 30 mM, pH 7.0). The mice were euthanized after 30 min of ATP injection and peritoneal cavity was lavaged with 5 ml PBS. Approximately 3.5 ml peritoneal lavage fluid was recovered from each mouse and centrifuged at 15 K rpm for 10 min at room temperature. The supernatant was subjected to IL-1β ELISA, using mouse IL-1β Quantikine ELISA kit (MLB00C, R&D systems) and following manufacturer's instructions.

***Mice RCT assays.*** WT C57BL6J mice were euthanized by CO2 inhalation and femoral bones were removed. The marrow was flushed out of the bones into a 50 ml sterile tube using a 10 ml syringe with a 26-gauge needle filled with sterile DMEM. Cells were centrifuged for 5 min at 1,800 rpm at 4°C, followed by two washes with sterile PBS. The cells were cultured for 11 to 14 days in DMEM supplemented with 20% L-cell conditioned medium, 10% fetal bovine serum and 1% penicillin/streptomycin. To generate foam cells, the tritium labeled $^3$H-cholesterol (2 µCi/ml; Perkin Elmer, Norwalk, USA) and 100 µg/ml acetylated LDL were mixed and incubated at 37°C. This cholesterol mixture was combined with DMEM containing 20% L-cell conditioned medium and BMDMs were incubated with cholesterol-labeled media for 48 h to generate foam cells. The cholesterol loaded foam cells were washed twice with DMEM prior to harvesting for *in vivo* injection, and ~2 million $^{3}$H-cholesterol dpm in a volume of 0.25 ml were transplanted subcutaneously on the upper back of recipient mice. At 24, 48, and 72 h post transplantation, plasma and feces samples were collected. Plasma radioactivity was determined, and total plasma dpm was calculated by estimating blood volume to be equal to 7% of the body weight and plasma to be 55% of the blood volume. RCT to the plasma was calculated as the % (dpm appearing in plasma/total dpm injected) of injected radioactivity. Collected feces were allowed to dry overnight at 55°C and then weighed. Feces were then hydrated in 10 ml of 50% ethanol solution followed by homogenization then an internal recovery standard of 10,000 dpm of $^{14}$C-cholesterol (Perkin-Elmer, Norwalk, USA) was added to each sample. The radioactivity was quantified as described in detail above. Upon sacrifice at 72 h, the liver was removed and weighed. A piece of liver, ~0.2 g, was isolated, weighed, suspended in PBS, homogenized, and a known amount of $^{14}$C-cholesterol radioactivity was added as a recovery standard. The radioactivity in an aliquot of 0.3 ml of the liver homogenate was measured by liquid scintillation counting The $^{14}$C-cholesterol dpm was used to back calculate the [$^{3}$H] recovery for the entire liver homogenate, which was further used to calculate the total amount of radioactivity in the liver. In addition to $^{14}$C-cholesterol internal standard, all RCT data was standardized to mouse body or fecal weight each harvest time point.

***Isolation of Bone marrow derived macrophages***: All experiments were performed in accordance with protocols approved by the Cleveland Clinic and Cleveland State University Institutional Animal Care and Use Committee (IACUC). WT C57BL6J mice were maintained on chow diet and sacrificed at 16 weeks of age. Femurs were collected to isolate and culture bone marrow macrophages using conditioned L-cell media. Mice were euthanized by $CO_2$ inhalation and femoral bones were removed. The marrow was flushed out of the bones into a 50 ml sterile tube using a 10 ml syringe with a 26-gauge needle filled with sterile DMEM. Cells were centrifuged for 5 min at 1,800 rpm at 4°C, followed by two washes with sterile PBS. The cells were resuspended in sterile-filtered BMDM growth media (DMEM with 7.6% fetal bovine serum, 15% L-cell conditioned media, and 0.76% penicillin/streptomycin mixture) and plated in 10 cm culture dishes and incubated at 37°C for 14 days. Cell media was replaced every 2–3 days for 2 weeks. The cells were routinely visualized under microscope for proliferation and differentiation into confluent BMDMs.

***Western blotting***: THP-1 macrophages were grown treated as indicated. The PBS-washed cell pellet was lysed in MPER lysis buffer containing protease inhibitors and PMSF. After discarding the nuclear pellet, the protein concentration was determined using the BCA protein assay (Pierce). 10-50 µg of cell protein samples were resolved on Novex 4-20% Tris- Glycine Gels (Invitrogen) and transferred onto polyvinylidene fluoride membranes (Invitrogen). Blots were incubated sequentially with 1:1000 rabbit polyclonal antibody against human IL-1β (Cell Signaling) and HRP-conjugated beta actin antibody (Sigma). The signal was detected with an enhanced chemiluminescent substrate (Pierce) and blots were imaged with iBright imaging system (ThermoFisher).

***Supplementary figures.***

***Fig. S1: Control mice injected with LPS or ATP alone, IL-1β levels in peritoneal lavage were determined by ELISA***.

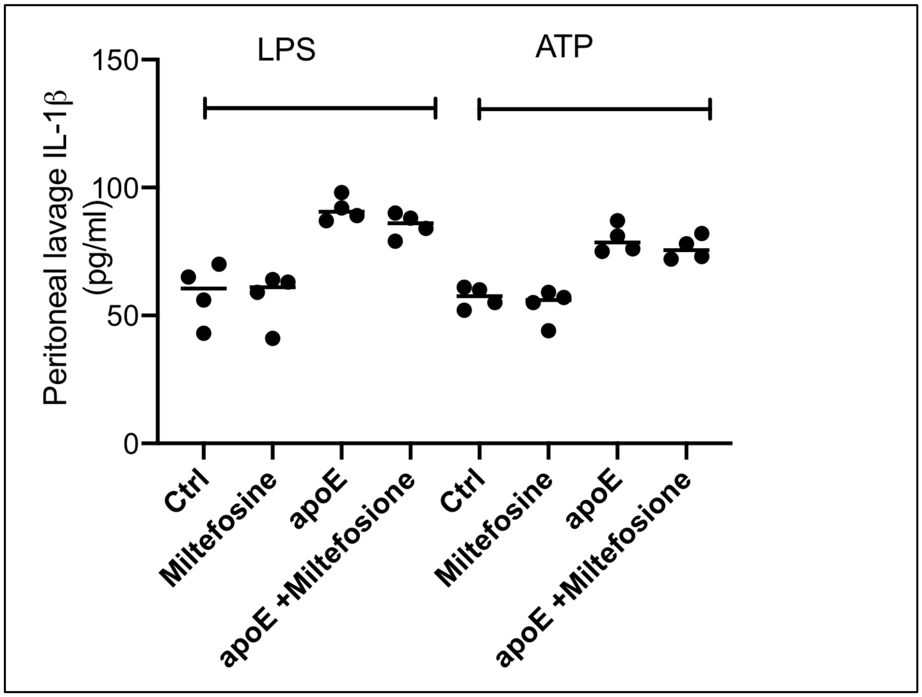

***Full images for western blot and microscopy images.***

***1) Full images for microscopy images shown in Fig. 1***

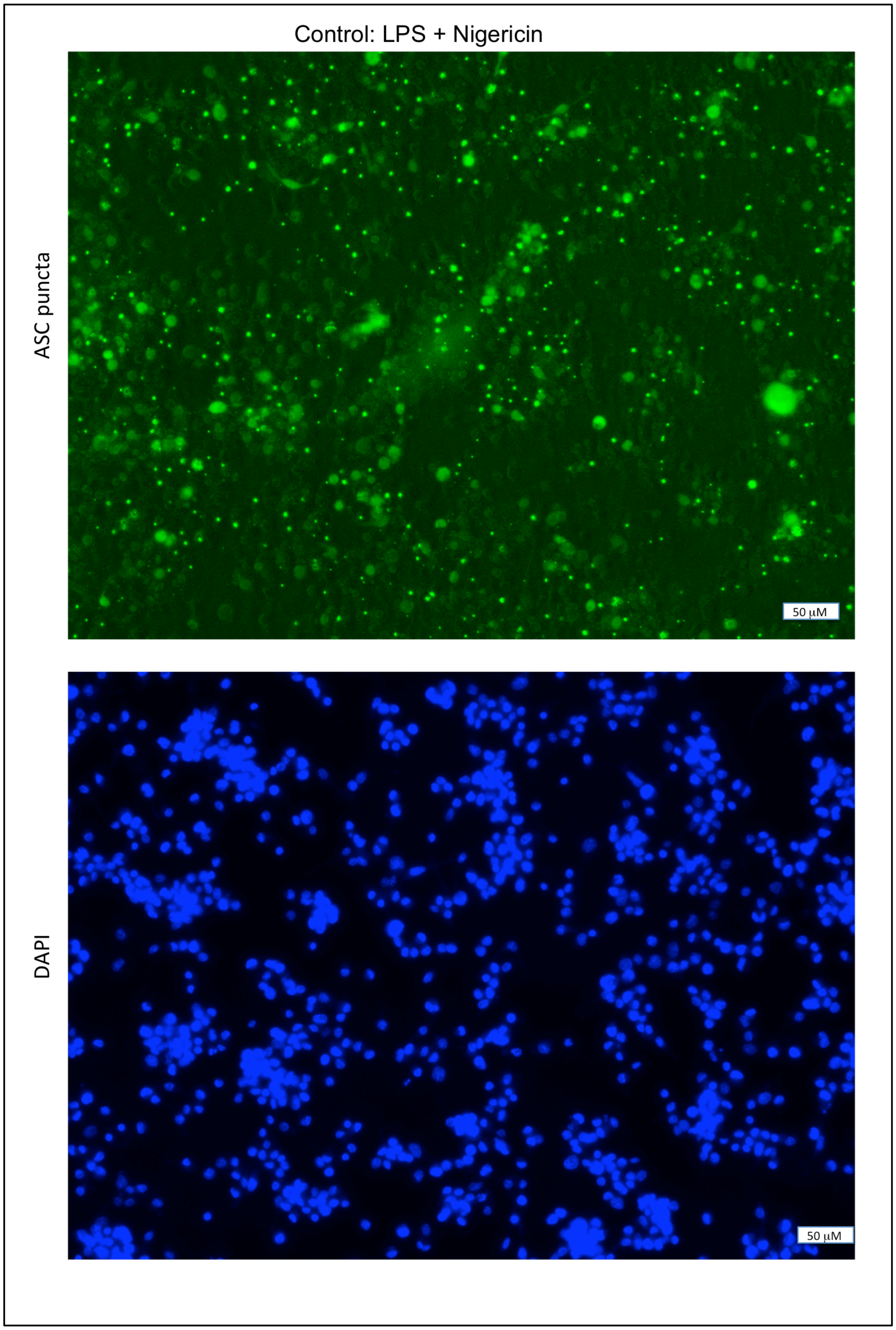

Miltefosine: LPS + Nigericin

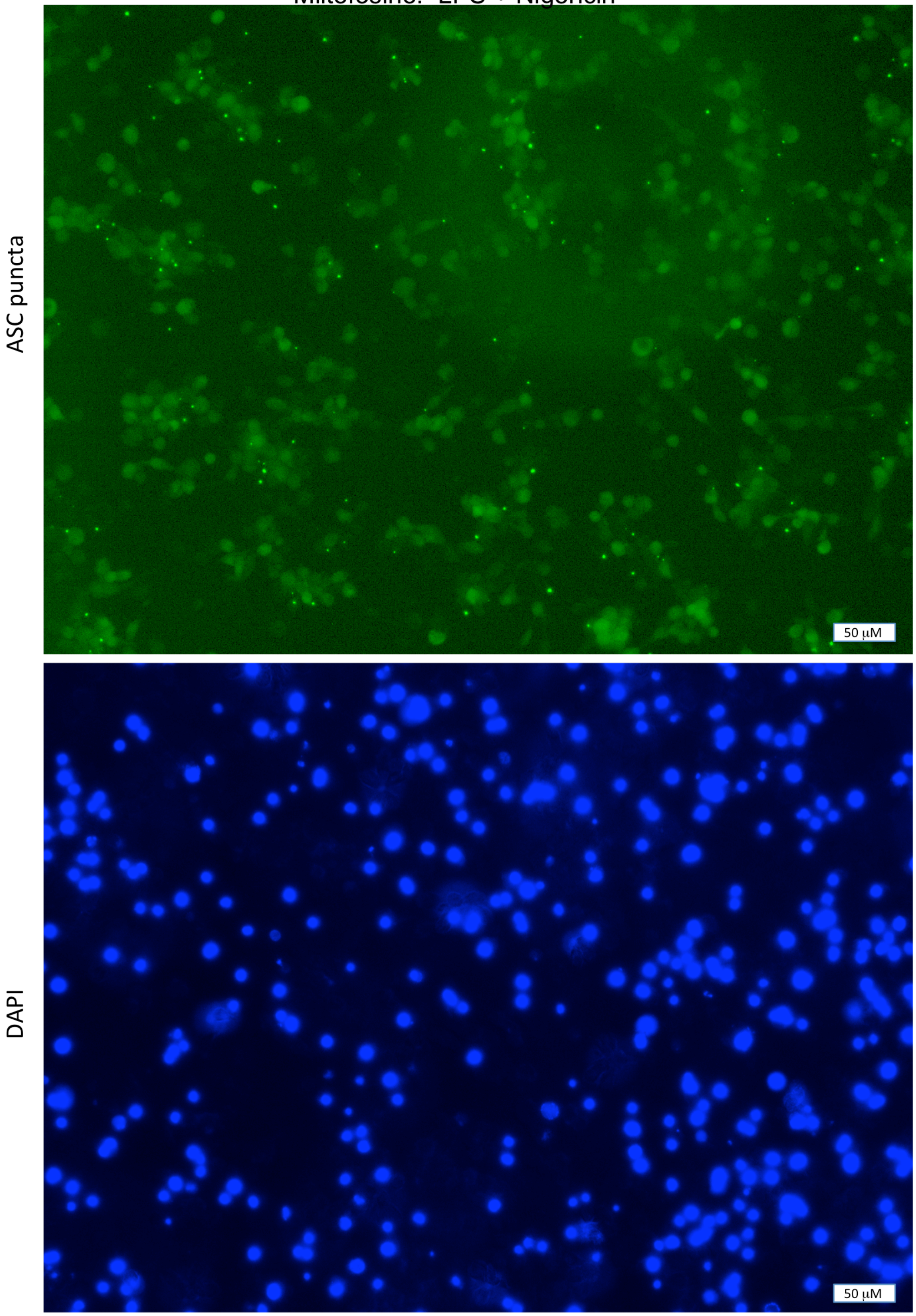

*2) Full images for western blot shown in Fig. 1*

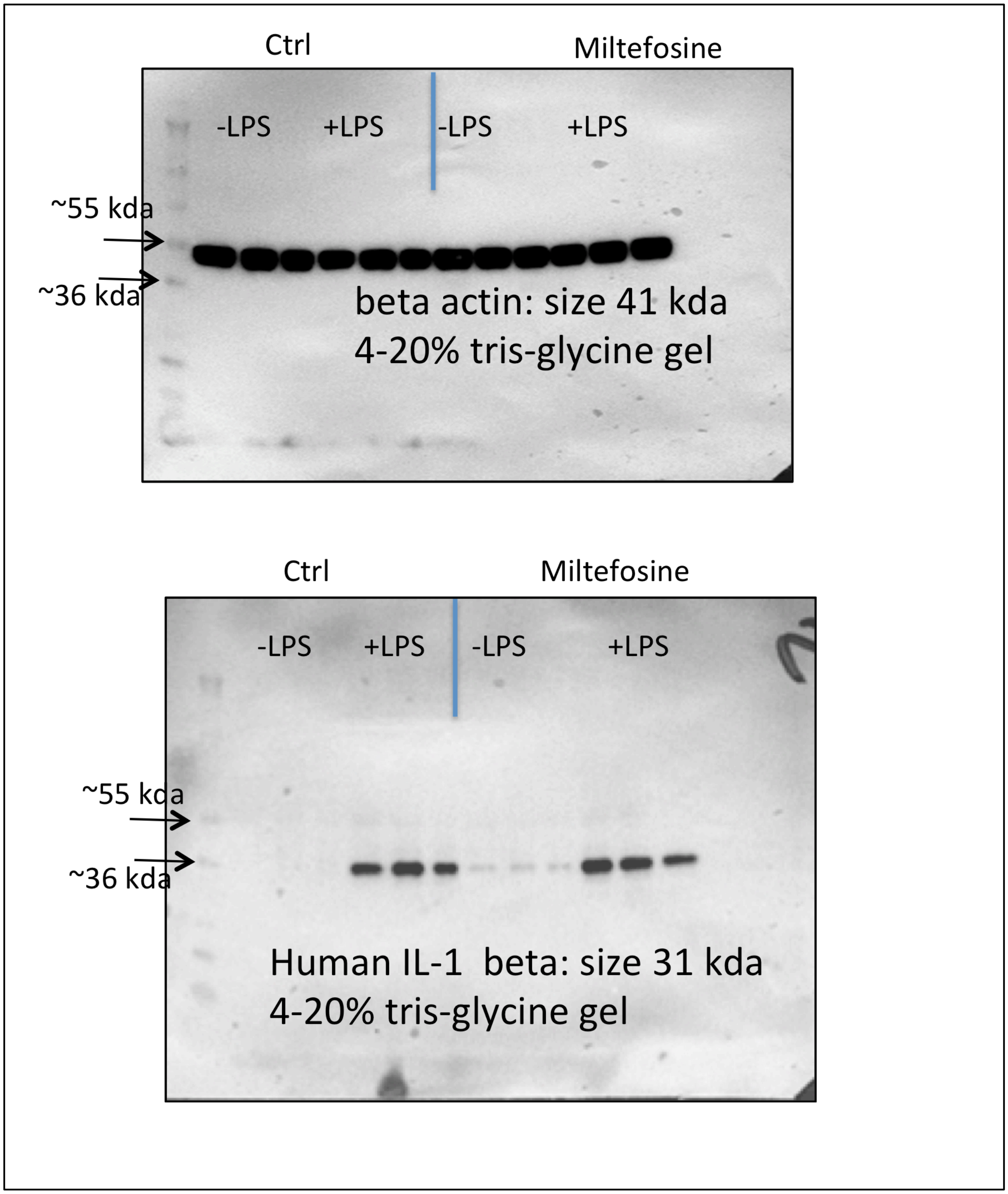
